# Neuronal activity induces myelin voltage changes that reflect action potential dependent myelin potassium buffering

**DOI:** 10.64898/2025.12.16.694668

**Authors:** Mélody Labarchède, Melina Petrel, Arne Battefeld

## Abstract

Vertebrate axons can be wrapped by myelin produced by oligodendrocytes. This cellular interaction ensures fast and accurate propagation of action potentials, but the physiology of the myelin sheath is almost completely unknown. To investigate the physiology of the myelin sheath, we implemented an imaging strategy that allowed optical measurements of myelin membrane voltage, with the aim to identify physiological changes of the myelin membrane during neuronal firing. We expressed the genetically encoded voltage indicator ASAP3 in mouse oligodendrocytes *in vivo* and subsequently investigated myelin physiology by optically measuring myelin membrane voltage. We found that myelin depolarizes during neuronal activity, which is blocked by inhibiting neuronal action potentials. Pharmacological and knock-out experiments of Kir4.1 showed that potassium uptake channels mediate action potential induced depolarization. Blocking myelin dependent potassium uptake and direct application of high potassium to identified axons induced axonal initiated and antidromic propagating action potentials. Our study shows that myelin is not an electrically passive insulator, but exhibits ion dynamics and its physiological response is fine tuned to neuronal activity. By facilitating potassium removal during action potentials, myelin supports high precision axonal firing. Genetically encoded sensors are thus a useful tool to study physiological properties of myelin, inaccessible by classical techniques.

**Highlights:**

- Optical imaging of myelin membrane potential
- Myelin sheaths exhibit depolarization in response to neuronal firing
- Depolarizations are partially mediated through Kir channels
- Potassium originates from axonal Kv channels

## Introduction

Myelin sheaths are highly compacted membranes around vertebrate axons. Myelin is considered isolating and an electrically passive structure that is required for fast saltatory action potential conduction. In myelinated axons, myelin contributes to organizing the spatial arrangement and clustering of axonal sodium and potassium ion channels. Although myelin plays a crucial role for axonal physiology, our knowledge of the physiological properties of myelin are very limited due to its small size making experimental approaches challenging.

Modelling predictions show that a small myelin conductance, up to 5 pS µm^-2^, could maintain saltatory conduction (Bakiri et al., 2011) predicting that open and functional channels can be found in the myelin sheath. Since this work, several ion channels and transporters have been found at the myelin sheath: The hyperpolarization-activated nucleotide gated channel HCN2 (Swire et al., 2021), the inward rectifying potassium channel Kir4.1 (Schirmer et al., 2018), the sodium-potassium co-transporter NKCC1 (Marshall-Phelps et al., 2020; Yamazaki et al., 2021) and previously the sodium-potassium ATPase (Zimmerman and Cammer, 1982). However, their impact on myelin physiology and the interplay with neuronal activity remains speculative and has not been directly demonstrated. On the axonal side, voltage gated potassium channels (Kv) are clustered in the juxta-paranodal region (Kv1) that is covered by the myelin sheath (Rasband and Shrager, 2000) and Kv7 channels cluster at the node of Ranvier (Pan et al., 2006) similar to the voltage gated sodium channel Nav1.6 (Caldwell et al., 2000).

Potassium (K^+^) homeostasis in the brain is highly regulated and alterations can lead to neuronal overexcitability and epilepsy (De Curtis et al., 2018). Oligodendrocyte cell bodies play a role in potassium clearing during neuronal stimulation via Kir4.1 channels and gap-junctions as previously demonstrated (Battefeld et al., 2016; Larson et al., 2018). At the myelin sheath our understanding of K^+^ regulation is limited. An ultrastructural study hypothesized that a pathway for potassium entry into the myelin sheath could be located in the juxta-paranodal myelin sheath, as connexin 29 hemichannels in myelin are facing axonal Kv1 channels (Rash et al., 2016). While connexin hemichannels are located within the inner most myelin layer facing the axon, Kir4.1 channels have also been shown to localize to the myelin sheath and paranodal loops (Kapell et al., 2023; Schirmer et al., 2018). We thus reasoned that membrane voltage changes of myelin in response to neuronal activity are highly probable as K^+^ homeostasis could be regulated on the level of myelin sheaths. However, somatic whole-cell recordings of oligodendrocyte are spatially limited due to the very low input resistance of oligodendrocytes (Battefeld et al., 2016) and direct electrical electrode recordings from the CNS myelin sheath are extremely challenging and limited to point recordings. To overcome this limitation we expressed the genetically encoded voltage indicator (GEVI) ASAP3 (Villette et al., 2019) in neocortical oligodendrocytes of mice and investigated myelin voltage at rest and in response to neuronal stimulation allowing novel insights into the physiology of the intricate myelin sheath structure and physiology.

## Results

To deliver and express ASAP3 in oligodendrocytes, we used an AAV based viral strategy allowing the transfection of mature oligodendrocytes (Hutson et al., 2012; Lawlor et al., 2009). We inserted ASAP3 into a viral backbone containing a 1.5 kb long myelin associated glycoprotein (*Mag*) promoter fragment that has been previously shown to be specific and effective in mature oligodendrocytes with little expression in other cells (von Jonquieres et al., 2016). The resulting ASAP3 construct (Figure 1A) was packaged into AAV2/8 and viral particles were injected by stereotaxic surgery into the deeper layers of the somato-sensory neocortex (Figure 1B). After 5 weeks of *in vivo* expression immunohistochemical analysis showed localization of ASAP3 in CNPase positive oligodendrocyte cell bodies and MBP or CNPase positive myelin sheaths (Figure 1C, Supplementary Figure 1A-B, n = 3 mice) and <10% expression in other brain cells (Supplementary Figure 1C, n = 12 sections from n = 3 mice). As ASAP3 is membrane anchored probe, we next asked in which myelin compartment ASAP3 is incorporated. We labelled the extracellular cpGFP domain of ASAP3 with immunogold particles and analyzed ASAP3 nanodomain localization in all myelin compartments (the inner tongue facing the axon, compact myelin and in the outer membrane layer) by transmission electron microscopy. On average 67% of ASAP3 positive gold particles were detected in the different myelin compartments (Figure 1D, E, Supplementary Figure 1D, n = 3 mice) with the majority of gold particles in compact myelin, demonstrating that ASAP3 expression leads to incorporation into all membranes of the myelin sheath. Based on the known dimensions of central myelin and the electron microscopy images we created a schematic working model of ASAP3 integration into the myelin sheath (Supplementary Figure 1E-G).

**Figure 1:**
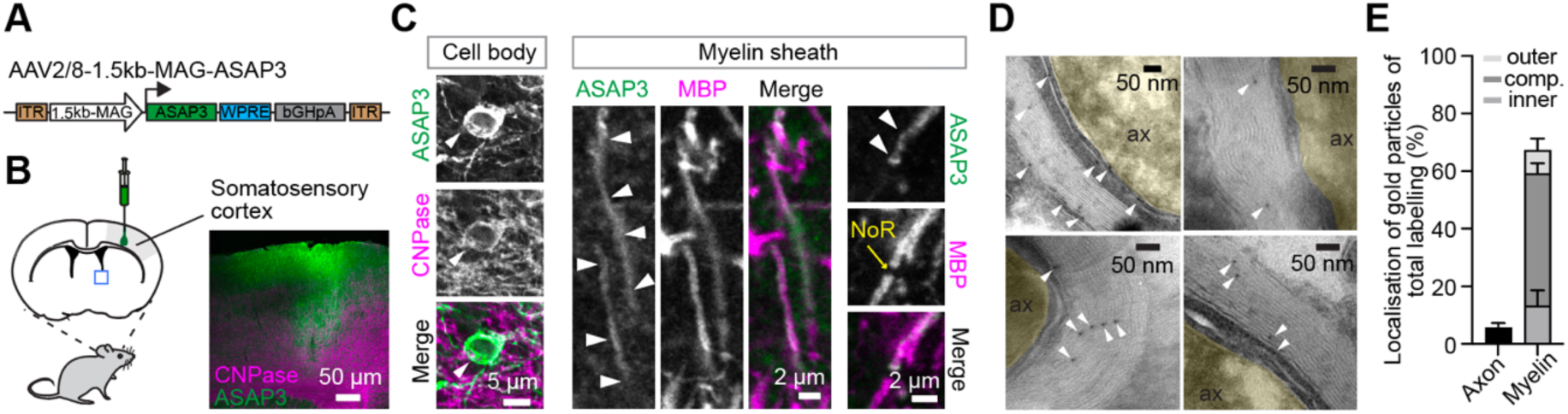
Viral mediated expression of the voltage sensor ASAP3 in mouse oligodendrocytes localizes ASAP3 to the myelin sheath. (A) AAV construct to target oligodendrocytes and express ASAP3 in vivo. ASAP3 expression is driven by a 1.5 kb fragment of the myelin-associated glycoprotein promoter. (B) Schematic of the AAV injection site in the somato-sensory neocortex and confocal example image of an injection site. (C) Confocal images of ASAP3 expression in oligodendrocyte cell bodies labeled with CNPase and myelin sheaths labelled with MBP. Arrowheads point to ASAP3 immunosignal at a cell body and in myelin sheaths of ASAP3 expressing oligodendrocytes thereby confirming that ASAP3 is located in all subcellular positions. ASAP3 immunosignal was amplified with an antibody against EGFP. (D) TEM images reveal the nanodomain localization of ASAP3 in the myelin sheath by immunogold labelling. We detected ASAP3 gold particles in all myelin compartments and very few in the adjacent axon (pseudocoloured in yellow). White arrowheads in all images point to 6 nm gold particles. (E) Quantification of immunogold particles shows that all myelin compartments (inner tongue, outer loops and compact myelin) are positive for ASAP3 gold particles (68 images from n = 3 mice), whereas labeling of the adjacent axon is very low.

Knowing that ASAP3 is well incorporated into myelin sheaths, we next investigated membrane voltage changes of myelin by optically recording ASAP3 fluorescence in acute slices of the somatosensory cortex. We targeted ASAP3 expressing oligodendrocytes in deeper cortical layers (Figure 2A) of young adult wild-type mice. Extracellular stimulation electrodes were placed in close proximity to the optical recording sites (average distance: cell body 26 ± 1.8 µm, n = 15; myelin 23.8 ±3.4 µm, n = 6 recordings). Stimulation resulted in a rapid depolarization of both: cell body and myelin sheaths, as measured by rapid imaging of ASAP3 (Figure 2B-D). The depolarization of cell bodies and myelin sheaths decayed exponentially and the decay time did not differ between these locations (Mann Whitney p = 0.73, n = 7 myelin sheaths, n = 6 cell bodies). The optically measured depolarization of the oligodendrocyte compartments was almost completely abolished after blocking action potentials with tetrodotoxin (TTX) or in absence of stimulation (Figure 2B,C, Supplementary Figure 2A), demonstrating that optically measured depolarization of oligodendrocyte cell bodies and myelin is linked to neuronal activity as previously shown for oligodendrocyte cell bodies (Battefeld et al., 2016; Larson et al., 2018). Changing the stimulation length from 0.1 s to 1 s with a constant stimulation frequency of 100 Hz resulted in depolarizations of soma and myelin for the duration of the stimulation time. Notably, 1-second-long stimulations did not linearly increase depolarization amplitude, presumably due to firing rate adaptation during these long depolarizations (Supplementary Figure 2B,C). The rise time of the depolarization was characterized by a fast voltage change at both cell body and myelin. In contrast, the voltage decay was prolonged for both subcellular locations (Figure 2D). In line, independent somatic whole-cell recordings combined with a single extracellular stimulation pulse led to a 3.2 ± 0.9 mV (n = 4) depolarization of oligodendrocytes that decayed slowly (tau decay 1.8 ± 0.6 s, n= 4, Supplementary Figure 2D,E). As extracellular stimulation and epifluorescence imaging can result in acquisition of signals that are out of the imaging plane, we additionally performed 2-photon imaging of single ASAP3 positive myelin sheaths (Figure 2E). As before, stimulation evoked an optically recorded voltage depolarization of myelin sheaths in response to extracellular stimulation (n = 4 sheaths, N = 3 slices from 2 mice). Together, our data reveal that the myelin sheath responds to neuronal firing with an increase in myelin conductance reflected in a myelin depolarization and reminiscent of an activation of ion channels in myelin.

**Figure 2:**
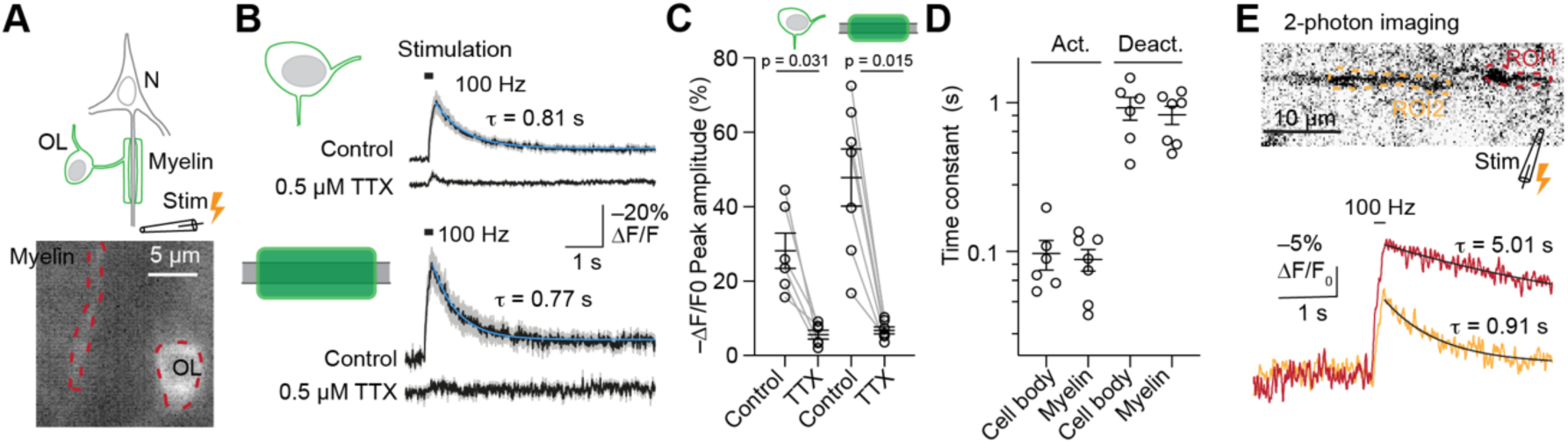
Optical recordings of oligodendrocyte membrane voltage reveal depolarization of myelin sheaths and cell bodies in response to neuronal stimulation. (A) Top: Experimental schematic of voltage imaging in oligodendrocytes. Bottom: ASAP3 fluorescence in an acute slice showing an oligodendrocyte cell body and myelin sheath. (B) Averages (black) and SEM (grey) of optically recorded voltage changes from oligodendrocyte cell body (n = 6) and myelin sheaths (n = 7 sheaths) in response to a 200 ms long electrical stimulation at 100 Hz. The decay of the voltage change was fit with a single exponential equation (blue line). 5 minutes of bath application of 0.5 µM TTX to block neuronal sodium channels strongly diminished the optically measured voltage change at the cell body and at myelin processes. (C) Summary data of peak fluorescence in control and after TTX application. The electrical stimulation dependent voltage change is almost completely abolished when blocking neuronal voltage dependent sodium channels. Cell body: n = 6 cells from 5 animals and 6 slices. Myelin: n = 7 sheaths from 6 slices and 5 animals. (D) Depolarization and decay time constants of the myelin and cell body voltage changes. N = 6 cell body, n = 7 myelin sheaths from 6 slices and 5 animals (E) Two-photon recording of ASAP-3 expressing myelin sheaths shows similar voltage responses after extracellular stimulation when compared to epifluorescence imaging. Two ROIs from two different myelin sheaths are shown.

What is the underlying ionic conductance that mediates myelin sheath depolarization? To investigate this question we targeted two channels that were previously shown to be located at the myelin sheath: HCN2 and Kir channels (Schirmer et al., 2018; Swire et al., 2021). Similar to the previous described experiments, we performed extracellular stimulation of myelin sheaths and subsequently washed in the HCN specific antagonist ZD7288 or the Kir channel blocker Barium (Ba^2+^) and repeated extracellular stimulation with same intensities (Figure 3A). Stimulation in the presence of 30 µM ZD7288 did not affect the amplitude of myelin depolarization or the fluorescence area (Figure 3B). Independent somatic whole-cell voltage recordings from grey matter oligodendrocyte somas showed a hyperpolarization of 3.8 ± 1.6 mV (n = 4) in current-clamp, confirming that somatic HCN currents are active at resting membrane potential in our experimental conditions, similar to published reports (Lyman et al., 2024; Swire et al., 2021). We next blocked Kir channels with 100 µM Ba^2+^. Comparing the control depolarization to the response in the presence of Ba^2+^ revealed a strong reduction of the peak amplitude and fluorescence area (Figure 3C). A likely candidate of the Kir family is the potassium uptake channel Kir4.1 (encoded by KCNJ10), which has been established to be located in compact myelin and paranodal loops by electron microscopy (Schirmer et al., 2018). Hence, we next addressed the specific involvement of Kir4.1 to myelin depolarizations. We generated an oligodendrocyte specific Kir4.1 knock-out by crossing inducible PLP-cre mice with floxed Kir4.1 (KCNJ10^flox/flox^) mice and expressed ASAP3 in the deeper layers of the cortex (Figure 3D) of these mice. Recordings of optical membrane voltage from myelin sheaths of Kir4.1 KO mice in response to electrical stimulation (Figure 3E) revealed a smaller peak amplitude and a reduced fluorescence area when compared to optical recordings from control littermates (Figure 3F). These experiments suggest that functional Kir4.1 channels remained or form stable heteromers with Kir5.1, which has been reported to be expressed in myelin on the mRNA level (Supplementary Figure 3A), but was not detected in top 1000 proteins under physiological conditions (Thakurela et al., 2016). In summary, these data demonstrate that increases in extracellular potassium concentration during neuronal activity result in a subsequent fast activation of Kir4.1 channels in myelin, reflected in myelin sheath depolarization. Myelin sheath voltage changes thus mirror the dynamic of extracellular potassium removal around axons.

**Figure 3:**
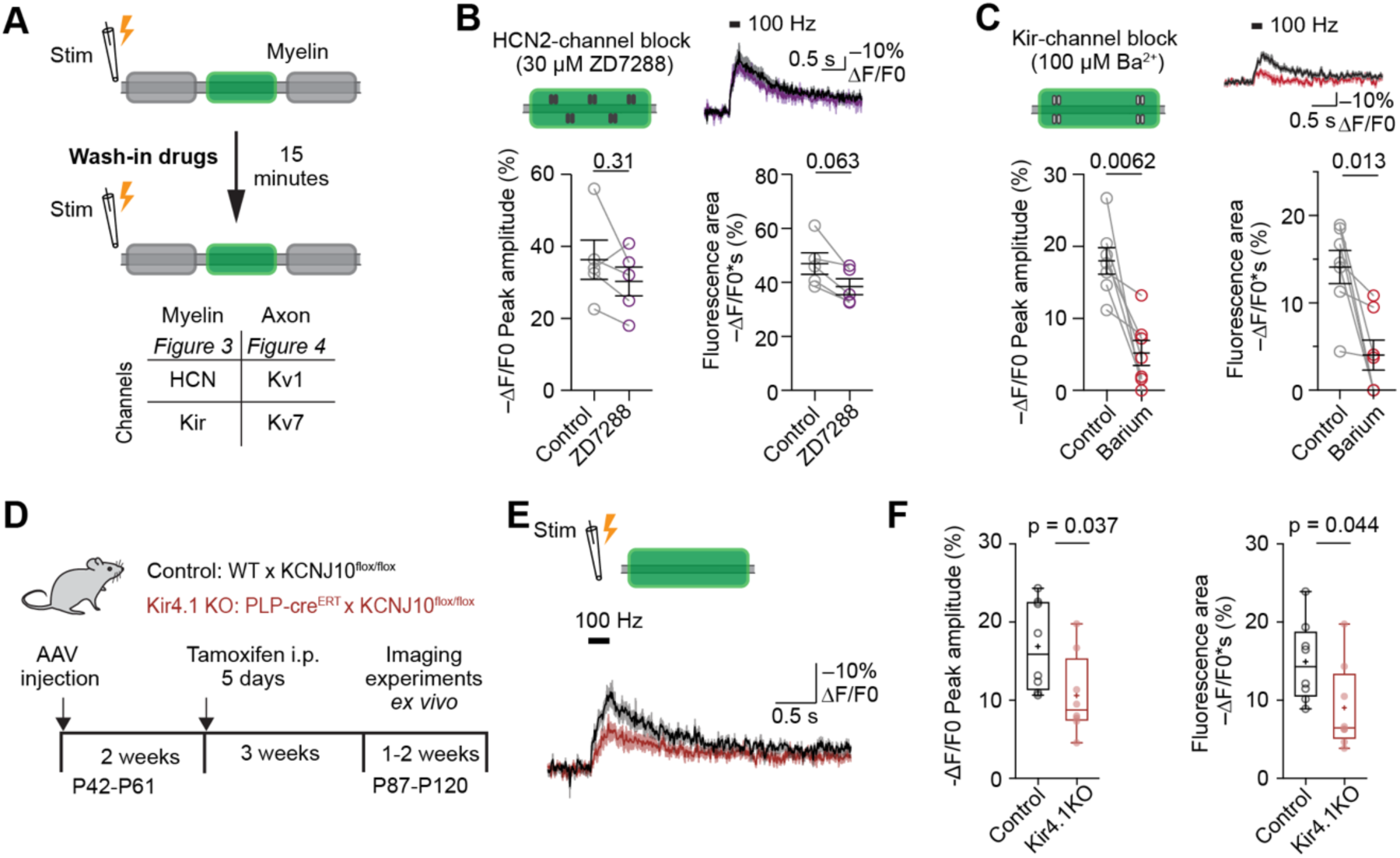
Kir4.1 channels contribute to optically measured myelin depolarizations. (A) Schematic of the experiment workflow. After identification of ASAP3 expressing myelin sheaths, axons were extracellularly stimulated and myelin voltage change was fluorescently recorded before and after application of drugs targeting known ion channels found in myelin or in the axonal nodal and juxtaparanodal compartment as shown in the table. (B) HCN2 channels located within in the myelin sheath were blocked with ZD7288. After 200 ms stimulation maximal optical voltage amplitude and area were unchanged by ZD7288. Wilcoxon test, 5 slices from 5 mice. (C) Application of Ba^2+^ to block Kir channels in myelin strongly reduced myelin depolarization as measured by amplitude and total fluorescence area. Wilcoxon test, 7 slices from 4 mice. (D) Schematic of the experimental timeline to target Kir4.1 (*Kcnj10*) deletion in mature oligodendrocytes and perform voltage imaging in Kir4.1 deleted animals. (E) Averaged optical myelin sheath voltage recordings and SEM from wild type (n = 8 slices) and Kir4.1^-/-^ (n = 8 slices) mice show a depolarization for both conditions after 200 ms stimulation. Kir4.1^-/-^ sheaths (red) show a reduced peak amplitude compared to control sheaths (black). (F) Quantifications of myelin depolarization of peak amplitude and the total fluorescence area are smaller in Kir4.1^-/-^ mice, which is consistent across several experiments when compared to control littermates. Control n = 8 slices from 3 mice; Kir4.1^-/-^ n = 8 slices from 5 mice.

Next, we assessed which neuronal ion channels contribute to myelin depolarization. Kv channels are activated during action potentials in cortical layer 5 neurons. In physiological conditions, axonal potassium Kv7 channels cluster at the nodes of Ranvier and Kv1 channels cluster under the juxtaparanodal loops (Rasband and Peles, 2021). We next assessed how potassium release from these channels impact myelin depolarization. We found that block of Kv1 channels, after bath application of 100 nM dendrotoxin-1, had little impact on amplitude and area of the optical measured myelin depolarization (Figure 4A). This result was surprising as we hypothesized an impact on the depolarization amplitude, but the access of dendrotoxin-1 to the juxtaparanodal Kv1 channels could have been limited. Blocking Kv7 channels with the specific blocker XE991 in the bath lead to a reduction in the depolarization amplitude and area, demonstrating the prolonged impact of Kv7 channel opening during action potentials (Figure 4B). As extracellular stimulation is not limited to a single axon, but optical recordings were performed from single myelin sheaths, other sources of potassium are presumably activated. In summary, our observations show that nodal potassium release depolarizes myelin sheath voltage.

**Figure 4:**
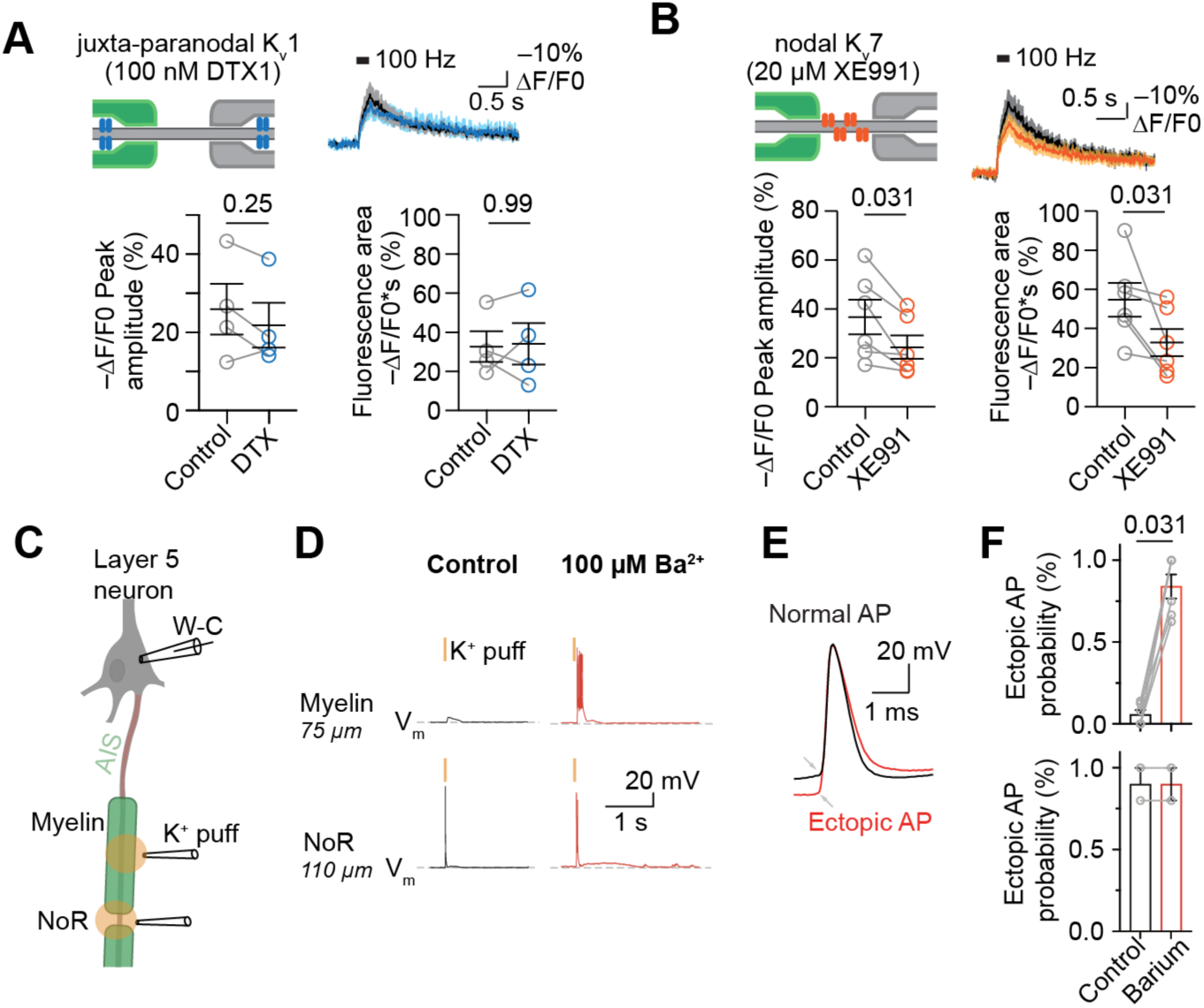
Blocking myelin Kir conductance evokes antidromic propagating ectopic action potentials. (A) Block of juxta-paranodal Kv1 channels with Dendrotoxin-1 (DTX1) does not impact the myelin voltage change. Wilcoxon test, 4 slices from 3 mice. (B) Blocking nodal Kv7 channels with XE991 reduces the peak myelin voltage and the fluorescence area suggesting that nodal potassium outflow contributes to myelin depolarization. Wilcoxon test, 6 slices from 5 mice. (C) Schematic of the axonal potassium application experiment. Layer 5 neurons were identified and high potassium solution was either applied to myelin or nodes to identify recording for axonal initiated spikes. (D) Somatic recordings in control and in the presence of 100 µM Ba^2+^ after a 50 ms potassium application to myelin or nodes of Ranvier show an increase in ectopically generated action potentials. (E) Example of a normal, AIS/somatic initiated AP, and an ectopic action potential that shows a about 10 mV more hyperpolarized threshold. (F) Quantification of ectopic AP recordings at the two puff locations myelin and node of Ranvier. Myelin n = 6 myelin sheath from 4 axons, Wilcoxon signed rank test. Node of Ranvier n = 2 nodes from 2 axons.

Finally, we addressed how local alterations of myelin sheath potassium buffering impact neuronal action potential generation at the axon. Previous work has demonstrated that demyelinated cortical layer 5 axons become more excitable and ectopic spikes (axonal) can be initiated (Hamada and Kole, 2015). To test whether the demonstrated potassium buffering capacity of myelin (Figure 2 and 3) contributes to the maintenance of axonal excitability under physiological conditions, we assessed the direct application of potassium to myelin sheaths while simultaneously recording electrically from layer 5 neurons (Figure 4C). We applied short puffs of high potassium solution either to myelin sheaths or the node of Ranvier from identified layer 5 neuron axons. When potassium was directly applied to the myelin sheath in control conditions, there was a low probability of ectopic action potentials. However, blocking Kir channels by application of 100 µM Ba^2+^ resulted in a strong increase of ectopic (antidromic) action potential probability (control: 6 ± 3%, Ba^2+^: 84 ± 18%, Figure 4D,E). At the node of Ranvier, potassium application did evoke ectopic action potentials in control and in the presence of Ba^2+^ (control: 90 ± 10%, Ba^2+^: 90 ± 10%, Figure 4D,E). These experiments show that the myelin sheath efficiently buffers local potassium increases and can shield the axon from potassium that is released from neighboring axons. Altering myelin buffering capacity, here by blocking Kir channels, can lead to axonal initiated spikes as a consequence. In summary, we demonstrate that during axonal action potentials, rapid inflow of potassium into myelin sheaths prevents high local potassium levels while at the same time restricting the generation of axonal ectopic action potentials probably by limiting potassium evoked axonal depolarization.

## Discussion

We optically measured voltage changes of myelin sheaths using a genetically encoded voltage indicator to address the physiological membrane voltage dynamics of the oligodendrocyte myelin sheath in the CNS. We found that myelin depolarizes rapidly in response to neuronal firing. Blocking action potentials with TTX abolished depolarization, we thus hypothesized that myelin sheath depolarizations are evoked by increases in extracellular potassium, released from axons during the repolarizing phase of the action potential. Pharmacological experiments revealed that barium strongly reduces the depolarization and in combination with Kir4.1 ablation experiments our results suggest that Kir4.1 mediated currents are activated by neuronal mediated extracellular K^+^ increase. Moreover, inhibiting neuronal potassium release from nodal Kv7 channels modulates the peak depolarization, further supporting that neuronal potassium release is a prerequisite for myelin sheath depolarization. Finally, direct application of K^+^ to myelin in the presence of barium leads to the occurrence of ectopic axonal action potentials suggesting that the myelin sheath protects from inaccurate axonal action potential generation. Our findings experimentally demonstrate that myelin is not only an insulating membrane, but myelin dynamically responds to changes in axonal activity and supports, via potassium buffering, high precision axonal firing. Moreover, these experiments link and establish that hypothesized K^+^ uptake pathways in myelin are functional, operate on the time scale of milliseconds and are putatively needed for long-term myelin and axon homeostasis (Rash, 2010; Schirmer et al., 2018).

What is the physiological functional relevance of myelin sheath depolarization? It has been demonstrated that extracellular potassium increases in white matter can act as signal to couple axonal energy demand to oligodendrocyte metabolic supply of axons (Looser et al., 2024). Our work directly demonstrates that myelin undergoes K^+^ mediated depolarization further supporting that ionic coupling between axon and the myelin sheath exists on a time scale that is locked to single action potential firing. While the work of Looser et al. (2024), links high firing rates with calcium increases, our previous experiments in mature grey matter myelin did not find increases of calcium in the myelin sheath during a natural firing stimulus (Battefeld et al., 2019). However, *in vivo*, optic nerve retinal ganglion cells fire at ∼35 Hz compared to a 1 to 5 Hz firing range of deeper layer neurons of the rodent somatosensory cortex (de Kock and Sakmann, 2009; Troy and Robson, 1992) thus potential differences exist between these two CNS locations.

Here, we observed reliable depolarizing responses in myelin that is abolished after blocking neuronal action potentials. A repetitive depolarizing response suggests that the underlying membrane potential of the myelin sheath is maintained by ion channels and pumps similar to the resting membrane potential of the oligodendrocyte cell body and other non-neuronal and neuronal cells. As hyperpolarization cyclic-nucleotide activated channels (HCN) are found in the myelin sheath membrane (Swire et al., 2021) the myelin sheath is probably more depolarized than a purely K^+^ permeable membrane. However, the nature of our imaging experiments did not allow the read-out of an absolute membrane voltage. Although we did not find that HCN2 blockage substantially altered myelin depolarization with optical membrane imaging in response to action potentials, we observed a resting membrane hyperpolarization in whole-cell patch-clamp recordings at oligodendrocyte somas. A depolarized myelin Vm is supported by measurements at the soma of grey and white matter oligodendrocytes when compared to astrocytes (Battefeld et al., 2016; Gipson and Bordey, 2002; Larson et al., 2018). cAMP plays a crucial role in shifting HCN2 voltage activation to more depolarized potentials (Moroni et al., 2000) making the cAMP pathway a powerful modulator of myelin sheath resting voltage. The g-protein coupled receptor GPR17 is implicated in reducing cAMP levels and its activation is linked to modulation of lactate levels in oligodendrocytes (Ou et al., 2019). However, GPR17 is only transiently expressed during oligodendrocyte maturation (Marques et al., 2016; Zhang et al., 2014), suggesting that at mature stages, as in our experiments, its influence on cAMP modulation is low. This is in-line with a depolarization of the resting membrane potential of oligodendrocytes with maturation (Battefeld et al., 2019) suggesting that GPR17 independent cAMP modulatory pathways or ionic mechanisms are responsible for resting membrane depolarization during oligodendrocyte maturation. Our experimental results provide a link between the maintenance of myelin membrane potential and neuronal K^+^ homeostasis to limit, on the one hand overexcitability and on the other hand regulate axon-myelin lactate metabolism (Looser et al., 2024).

Early work on axonal physiology revealed that prolonged firing releases potassium into the extracellular space (Frankenhaeuser and Hodgkin, 1956). In the myelinated axon, K^+^ diffusion into oligodendrocytes is an efficient way of clearing up extracellular K^+^ and limiting the depolarizing impact on the neuronal resting potential and therefore increased (non-synchronized) firing as well as limiting a smaller action potential amplitude due to reduced availability of sodium channels. Based on our experimental data we suggest that the myelin sheath carries out potassium buffering function and thus oligodendrocytes are the primary cells that support potassium removal and buffering of the myelinated axon as they are in a privileged anatomical location. It is notable, that depolarization of the myelin sheath was not localized and occurred throughout one sheath. This could be due to technical aspects as a low imaging frequency (∼200 Hz) or suggests that K^+^ can come from several sources. In support of the latter argument, we could not identify local depolarization differences in one imaged sheath.

Our pharmacological experiments did not show any change of Kv1 channel mediated potassium outflow, which could be due to low diffusion accessibility of dendrotoxin-1 to the myelin covered channels. Hence, we can not clearly conclude the contribution of Kv1 channels that are located in the paranodal region under the myelin sheath and their impact on myelin depolarization. However, as we demonstrate K^+^ buffering of myelin, our experiments still suggest that Kv1 released K^+^ can be directly taken up by the myelin sheath, but a clear experimental demonstration lacks. Future experiments, could thus further investigate the sub-myelin Kv1 contribution to myelin depolarization which could include faster optical recordings in Kv1 mutant mice, mathematical modelling and more precise activation of the axon-myelin unit by combining optogenetic neuronal activation and voltage imaging. In addition, potassium flow might be directly monitored for instance by utilizing improved genetically encoded sensors for potassium (Torres Cabán et al., 2022) that could be expressed with our strategy in oligodendrocytes.

Depolarization of myelin points to a mechanism that couples neuronal and oligodendrocyte physiology potentially providing a read-out of neuronal activity on a millisecond time scale at spatial scale of single myelin sheaths. It remains unaddressed how K^+^ dynamics are within the multi-membrane myelin layer. Due to its size we can only speculate on the mechanism. Gap junctions that connect the different myelin layers can mediate K^+^ flow within the myelin (Orthmann-Murphy et al., 2007). Similarly, claudin-11 tight junctions in myelin could contribute to current flow across the myelin sheath (Devaux and Gow, 2008). It remains to be demonstrated whether K^+^ can flow across the myelin layers or whether K^+^ uptake on the inner cytoplasmic tongue only contributes to a gradient needed to drive the Na^+^K^+^-ATPase as shown for tight junctions in epithelial tissues (Rajasekaran et al., 2007).

K^+^ concentrations in the CNS are tightly controlled by several redundant mechanisms demonstrating the importance of K^+^ regulation. Proper ion homeostasis and especially potassium homeostasis, has been linked to myelin maintenance. For instance, the loss of NKCC1 in oligodendrocytes disrupting the co-transport of sodium, potassium and chloride (Marshall-Phelps et al., 2020). Indirect evidence suggests that MLC1 dysfunction in astrocytes which regulates potassium and water homeostasis (Min and van der Knaap, 2018) can lead to myelin vacuolization and alterations of myelin integrity. Although previously suggested that Kir4.1 channels mediate axonal potassium uptake and its ablation leads to long-term axonal alterations (Schirmer et al., 2018), we here demonstrate directly that axonal potassium outflow leads to myelin depolarization and potassium uptake via Kir4.1 located in myelin. Recent work in the EAE model suggests that the Kv7 channel opener retigabine can improve clinical signs in the absence Kir4.1 in oligodendrocytes (Kapell et al., 2023). These latter results appear contradictory to our findings as more potassium outflow would lead to stronger myelin depolarization and put more pressure on the buffering capacity of myelin. However, our work focuses on fast dynamic myelin physiology whereas in the case of longer-term pharmacological treatments other homeostatic changes could appear. Supporting our data is recent work, that has shown that changes in extracellular potassium homeostasis are found in several neurodegenerative disease models (Ding et al., 2024) which have been associated with alterations or loss of oligodendrocytes (Kang et al., 2013; Lim et al., 2022; Sasmita et al., 2024; Schneeberger et al., 2025). Future investigations could address the role of oligodendrocyte and myelin potassium buffering in these neurodegenerative models.

Experiments were performed in cortical oligodendrocytes and we cannot exclude that in other brain regions myelin responses to neuronal activity are different. Myelinating oligodendrocytes downregulate sodium channels, however, voltage gated calcium channels have been previously reported (Paez and Lyons, 2020). With our imaging approach, millisecond fast transient depolarizations of voltage-gated calcium channels (Chemin et al., 2002) cannot be monitored, but after the initial depolarization, slower deactivation kinetics could be partially contributing during the return of myelin membrane voltage to resting levels. However, expression of calcium channels in cortical oligodendrocytes – MOL type 5 and 6 (Marques et al., 2016) are in majority high-voltage activated channels requiring strong depolarization of the myelin membrane. Although the calibration of the optical signal with voltage change in myelin is technically challenging, in previous work no action potential linked calcium increases were reported in grey matter myelin (Battefeld et al., 2019) indirectly showing the absence of voltage-gated calcium currents during physiological stimulation paradigms in cortical oligodendrocytes. In contrast, myelin in the optic nerve has recently been shown to respond with calcium increase in response to 30-second-long stimulations (Looser et al., 2024) and differences between white and grey matter oligodendrocytes could exist.

Our data demonstrates that the myelin sheath is not a passive compartment, but myelin conductance is interlinked with neuronal activity reflected in myelin depolarization during action potential firing. These results show that myelin physiology can be explored by applying novel tools. In the future this approach opens up further possibilities to study myelin physiology by focusing on novel targets that can be further explored.

## Materials and Methods

### Data Availability statement

All data generated or analyzed during this study are included in the manuscript and in the associated supplemental files.

### Ethical approval

All experimental procedures were in agreement with EU directive 2010/63/EU, validated by the local ethics committee (CEEA-50 Bordeaux) and approved by the French Ministry of Higher Education, Research and Innovation for handling animals (APAFIS #27726). To reduce the overall number of mice, we used one brain hemisphere for physiology experiments and the other hemisphere for histology, when applicable.

### Animals

For all ex vivo and histology experiments we used male and female mice with an age between 12 to 22 weeks. Mice were bred in an access restricted specific pathogen free facility and then transferred to an experimental facility for all further experiments. Mice were housed in individually ventilated cages in collectivity with a reverse 12 h day/12 h night cycle (light on at 7 am, light off at 7 pm) with constant temperature and humidity and water and food ad libitum. For the described experiments we used either C57Bl6/J or transgenic lines that were maintained on a C57Bl6/J background. For the majority of experiments we used C57Bl6/J mice, wild-type littermates from a Cnp-mEGFP reporter strain or mice expressing mEGFP (Deng et al., 2014). For subsets of experiments we crossed Plp1creER mice (B6.Cg-Tg(Plp1-cre/ERT)3Pop/J (Doerflinger et al., 2003)) with a floxed tdTomato mouse-line Ai9T (Gt(Rosa)26Sortm6(CAG-tdTomato)Hze, (Madisen et al., 2010). For 2P experiments we used PV-cre mice (Hippenmeyer et al., 2005) crossed with Ai9t reporter mice. We additionally crossed PLP1creERxAi9T with Kir4.1fl/fl mice (Djukic et al., 2007) to induce an oligodendrocytic specific knock-out of Kir4.1 (KCNJ10). When applicable, generated mice that were cre negative (cre^-/-^) were used as controls. Cre expression and recombination of all crosses involving PLP1-creER mice was induced by intra-peritoneal injections of tamoxifen for 5 consecutive days (for more details see **Tamoxifen preparation and injection).**

### Genotyping

Animal genotyping was performed by the central genotyping facility (Neurocentre Magendie). The following primers for genotyping were used:

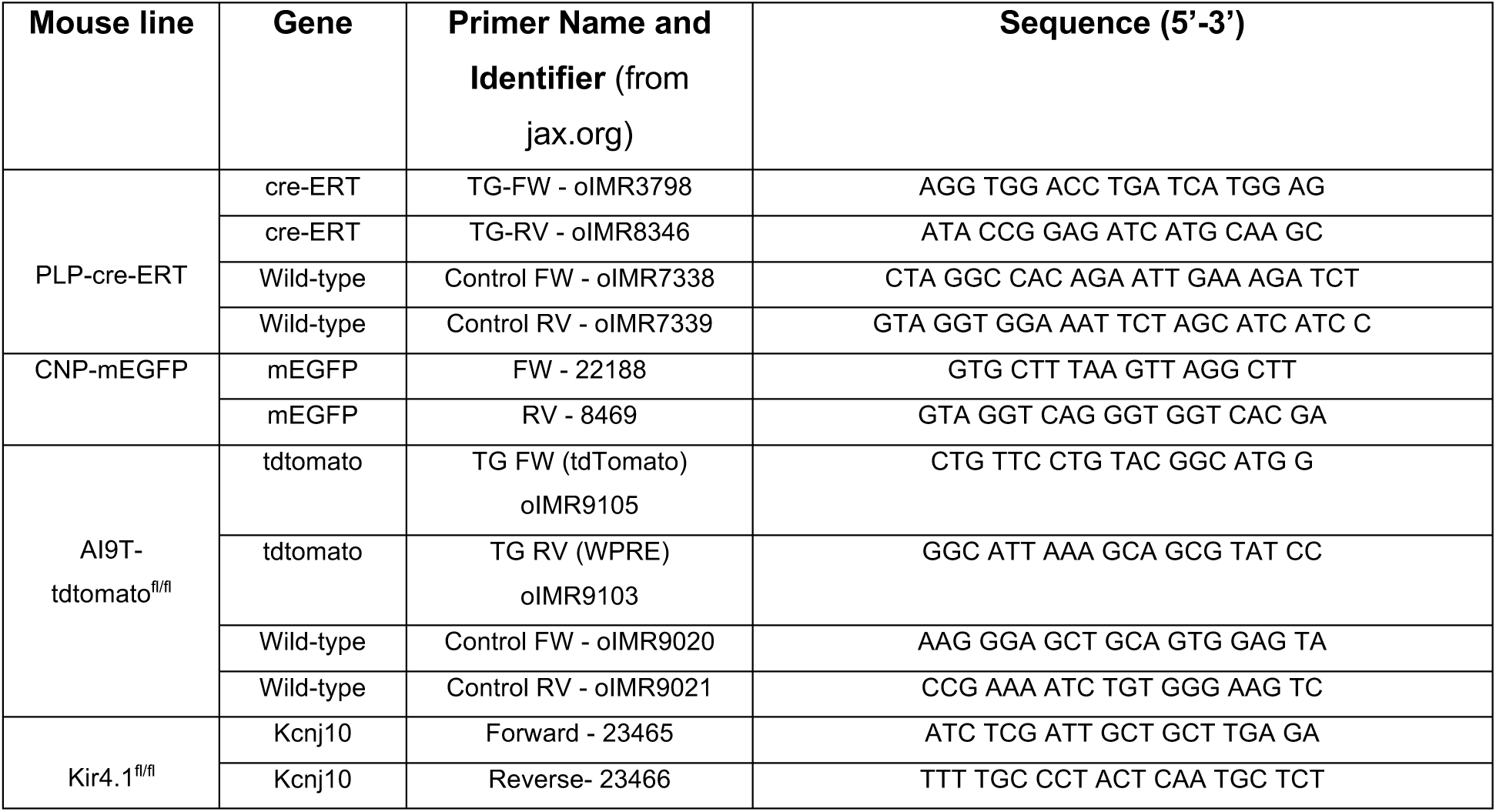

### Cloning of AAV constructs

ASAP3 (RRID:Addgene_132331) was synthesized (GeneUniversal Inc., Newark, DE, USA) flanked by a 5’ XbaI and 3’ BamHI restriction site. After restriction enzyme digestion with XbaI and BamHI (both Fast Digest, Thermo Scientific, USA) the ASAP3 ORF was ligated into an oligodendrocyte specific adeno-associated virus (AAV) plasmid, in which expression was driven by a myelin-associated glycoprotein (MAG) promoter of 1.5 kb length (von Jonquieres et al., 2016). The original 1.5 kb MAG promoter plasmid expressing GFP was modified to facilitate cloning into the AAV plasmid. A short multi-cloning site with the sequence 5’ AGATCTACGCGTGAATTCTAGATATCATATGGATCCTGCAG 3’, was inserted between the MAG promoter and the WPRE sequence and the GFP open-reading frame was removed (pAAV-1.5MAG-MCS-WPRE – Addgene #196547). This backbone vector was then cut with XbaI and BamHI and the ASAP3 ORF was ligated (T4 DNA Ligase, #15224017, Invitrogen) into the empty plasmid and designated pAAV-1.5MAG-ASAP3-WPRE (Addgene #196548)). We verified all sequence inserts by Sanger sequencing. During cloning, plasmids were amplified using NEB stable competent bacteria (C3040, NEB, MA, USA) and grown at 30°C to reduce recombination of the AAV ITRs. After producing larger amounts of plasmid DNA, the percentage of ITR recombination was checked with Sma1 digestion and was below 10%. Both plasmids were deposited at Addgene.

### AAV production

Recombinant AAV2/8 vectors (rAAV2/8) were produced at the in-house viral core facility. Low passage HEK-293T/17 cells (ATCC; cat number CRL-11268) were maintained in DMEM at 37°C and 5% CO2. HEK cells were triple transfected with plasmids for ASAP3 (this study), AAV *rep* and *cap* genes (pAAV2/8 was a gift from James M. Wilson; Addgene plasmid # 112864; RRID:Addgene_112864), and adenoviral helper functions (pAdDeltaF6 was a gift from James M. Wilson; Addgene plasmid # 112867; RRID:Addgene_112867) using polyethylenimine (PEI). HEK cells were harvested 72 hours post transfection and resuspended in lysis buffer (150 mM NaCl, 50 mM Tris-HCl pH 8.5) and lysed by 3 freeze-thaw cycles (37°C/–80°C). The cell lysate was treated with 150 U/ml benzonase (Sigma-Aldrich, Saint-Quentin-Fallavier Cedex, France) for 1 hour at 37°C and the crude lysate was clarified by centrifugation. Subsequently, rAAVs were purified by iodixanol step gradient centrifugation, concentrated, and buffer-exchanged into lactated Ringer’s solution (Baxter, Deerfield, IL, USA) using vivaspin20 100 kDa cut-off concentrator (Sartorius Stedim, Göttingen, Germany). The genome-containing particle (gcp) titre was determined by quantitative real-time PCR using SYBR green master mix (04887352001, Roche Diagnostics, Meylan, France) with primers specific for the AAV2 ITRs (fwd 5′-GGAACCCCTAGTGATGGAGTT-3′; rev 5′-CGGCCTCAGTGAGCGA-3′) on a Light Cycler 480 instrument. Concentrated virus was stored in aliquots at –80°C. The titre for the prepared AAV was determined to be 2.6x10E^13^ and the virus was used undiluted for stereotaxic injections.

### Stereotaxic surgeries

For stereotaxic surgeries we used mice of the following lines: C57Bl6/J, PLP1-crexAi9T, PLP1-cre^+^::Kir4.1^fl/fl^, PLP1-cre^-/-^::Kir4.1^fl/fl^. Anesthesia was induced with 4% isoflurane (Iso-vet, Piramal Healthcare, Bombay, India) and maintained at 1.5% using a mask fitted on the snout during duration of the surgery. For relieving pain, mice received an intra-peritoneal injection of buprenorphine (buprecare, 0,1 mg/kg) and carprofene (rimadyl, 5 mg/kg) before and after surgery. Mice were fixed into a stereotaxic frame (RWD Lifesciences, Mainz, Germany) and placed on a heating pad (set to 38°C) to maintain the body temperature during the surgery. To prevent drying-out of eyes, an ophthalmic gel (ocry-gel, TVM lab, Lempdes, France) was put on each eye and regularly refreshed during the surgery. A sub-cutaneous injection of lidocaine (Lurocaine, 7mg/kg) was made at the incision site for local anesthesia. After the skull bone was exposed the somatosensory cortex injection coordinates (from Bregma: ML: +/- 1.8 mm, AP: 0.2 mm and -0.6 mm, DV: -0.7 mm) were determined bilaterally and at these locations a small hole was carefully drilled. Subsequently the tip of the glass injection pipette (IntraMARK Brand, ref: 708707, Sigma-Aldrich) was lowered to the injection coordinate and the adeno-associated virus was injected using a picospritzer (Picopump PV820-A, World Precision Instruments, Sarasota, USA) connected to the injection pipette. At each injection site we applied a volume of 100 nL. Micropipettes sized between 30 and 60 µm were front filled before the experiments using negative pressure and kept at 4°C during the surgery. The skin was sutured with non-resorbable threads (EH7471H, Ethicon, Johnson and Johnson, France) and mice were monitored and kept on a heating pad until their recovery from anaesthesia (usually around 30 minutes). At the end animals were placed back in their home-cage. Post-operative check was performed for the following three days and included weight charting and behavioral assessment parameters. When sutures remained, they were removed 1 week after the surgery.

### Tamoxifen preparation and injection

A tamoxifen (T5648, Sigma-Aldrich) stock solution of 20 mg/ml was prepared in sterile filtered corn oil (C8267, Sigma-Aldrich) and stored in aliquots at –20°C. Once thawed, an aliquot was kept at 4°C and was used within one week. One week after the stereotaxic surgery, mice received intraperitoneal injections of tamoxifen (75mg/kg) for five consecutive days.

### Preparation of acute brain slices

Mice were deeply anesthetized with an intraperitoneal injection of a mixture of ketamine (100 mg/kg) and xylazine (20 mg/kg) diluted in 0.9% NaCl. After absence of reflexes (tail and toe) the thorax of mice was carefully opened and the mouse was transcardially perfused with about 10 ml of ice cold modified artificial cerebrospinal fluid (ACSF) composed of (in mM) 60 NaCl, 25 NaHCO3, 1.25 NaH2PO4, 2.5 KCl, 100 sucrose, 1 CaCl2, 5 MgCl2 and 20 glucose that was saturated with a mixture of 95% O2 and 5% CO2. After the blood was completely removed, mice were decapitated, the brain dissected and the hemispheres separated. In some experiments, one hemisphere was quickly put into ice-cold modified ACSF while the other was drop-fixed in 4% paraformaldehyde (Sigma-Aldrich) for 2 hours at room temperature and subsequently used for histology. The hemisphere in ice-cold ACSF was removed and glued to a specimen holder. A piece of agarose gel (1% prepared in ACSF) was glued behind the hemisphere for more stability during slice preparation. The tissue was then covered with ice-cold ACSF and mounted on a vibratome stage (VT1200S, Leica, Germany). We prepared parasagittal and coronal slices of 300 µm thickness that contained the injection site. Slices were collected and kept for 45 min at 37°C in carbogen saturated storage ACSF composed of (in mM) 125 NaCl, 25 NaHCO3, 1.25 NaH2PO4, 3 KCl, 1 CaCl2, 6 MgCl2 and 20 glucose and kept in this solution for the rest of the experimental day at room temperature.

### Epifluorescence voltage imaging

A slice was transferred to a motorized upright microscope (LN scope, Luigs-Neumann, Ratingen, Germany) equipped with Olympus optical elements and a submerged microscope chamber. The slice was continuously perfused with warmed (32 ± 1°C) recording ACSF composed of (in mM) 125 NaCl, 25 NaHCO3, 1.25 NaH2PO4, 3 KCl, 2 CaCl2, 1 MgCl2 and 25 glucose. Identification of cellular structures was performed with infra-red illumination (740 nm LED, LED engine/Osram, München, Germany) and an oblique contrast condenser. ASAP3 was excited by a single collimated LED that was mounted on a large passive heatsink with a peak emission of 470-472 nm (SP-01-B6, Luxeon Star Rebel LED, Lethbridge, Alberta, Canada). The epifluorescence excitation light was bandpass filtered (ET470/24 nm, Chroma, Bellow Falls, VT, USA) and then either directly coupled into the light path or reflected off a dichroic mirror (FF573-Di01, Semrock, IDEX Health & Science, LLC, Rochester, NY, USA) before the light was reflected onto the sample with a dichroic mirror (Di02-R-488, Semrock) mounted in a dual infinity cube (Cairn-Research, Faversham, Kent, UK). The emission light was reflected off a dichroic mirror (FF685-Di02, Semrock,) into an OptoSplit III (Cairn-Research) where the light was either reflected off a mirror or passed through passed through a triple dichroic mirror (69008bs, Chroma). ASAP3 emission light was bandpass emission filtered (ET535/30m, Chroma) and in case of the triple dichroic mirror the longer wavelength light was left unfiltered. Images were focused onto a sCMOS camera (95B, Teledyne Photometrics, Tucson, AZ, USA) either as a single image or side by side. A 63× 1.0 NA water immersion objective (Olympus) was used for all epifluorescence imaging experiments. Oligodendrocytes and myelin sheaths were identified by their expression of ASAP3, their soma morphology and appearance in oblique contrast and their location within the tissue. The camera exposure time (3 ms) and acquisition frequency was set in Micro-manager that was also used to acquire images and store images (Edelstein et al., 2014). For time-series images we used the multiple-acquisition module and an imaging frequency of 200 Hz with a reduced ROI. The camera and stimulation electrode were triggered by SutterPatch software and the 95B trigger-out was fed into a cyclops LED driver (Open-Ephys, (Newman et al., 2015)) controlling the imaging LED, leading to an effective global shutter of each image.

### 2-photon imaging

For 2-photon imaging, a motorized upright microscope (FEMTO3D-RC, Femtonics, Budapest, Hungary) with a fixed stage was used located at the central imaging facility. The microscope was equipped with a Nikon 25× NA 1.10 objective (ApoLWD). For 2-photon excitation, a femtosecond tunable laser was used (Chameleon Vision 2, Coherent, Santa Clara, CA, USA) and laser power was modulated by a Pockels cell (Conoptics, Danbury, CT, USA). Fast imaging was achieved with a galvo scanner and fluorescence signals were collected on NDD GaAsP photomultiplier tubes equipped with a green bandpass filter (490 to 550 nm, Chroma). Line scans were performed on myelin sheaths with the laser wavelength tuned to 920 nm and the laser power set between 1 to 2%.

### Electrical stimulation

For electrical stimulation, borosilicate glass capillaries (BF150-86-10, Sutter Instrument, Novato, CA, USA) were pulled with a puller (Sutter Instrument, Model P-97) to the shape of patch-pipettes and filled with ACSF. A sliver chloride wire was inserted into the pipette and the reference silver wire was wrapped around the stimulation pipette ending close to the pipette tip. Subsequently, the stimulation pipette was placed close to the recording site and pulses of 1 mA were applied using a stimulus isolator (Iso-Flex, A.M.P.I., Jerusalem, Israel). When trains of stimulations were applied these lasted from 100 to 1000 ms depending on the experiment and frequencies between 50 to 100 Hz. Timing of electrical stimulation was controlled by SutterPatch software (IPA2, Sutter Instruments, Novato, CA, USA) and synchronized to image acquisition. For 2-photon experiments, extracellular stimulation intensity was set to 0.4 mA and TTL pulses were given at 100 Hz intervals for 100 ms (MES software, Femtonics). For all experiments the duration of stimulation pulses was set to 100 µs.

### Whole-cell patch-clamp recordings

Slices were put under an upright microscope in a submerged chamber and continuously perfused with ACSF (see Epifluorescence voltage imaging). Whole-cell recordings were established with glass pipettes of 6-7 MΩ resistance that were backfilled filled with intracellular solution (in mM) 130 K-gluconate, 10 KCl, 10 HEPES, 4 Mg2-ATP, 0.3 Na-GTP, 10 Na2-phosphocreatine, 15 mM biocytin and a pH of 7.25. All recordings were performed with a Sutter double IPA amplifier (Sutter Instruments) that was software controlled by SutterPatch software. Membrane voltage responses were recorded in current-clamp and sampled with a minimum frequency of 20 kHz. Full compensation of bridge balance and capacitance was applied and checked before the start of a recording. Oligodendrocytes were identified in CNP-mGFP positive mice in the somatosensory cortex and whole cell-recordings were established at their cell body. Current and voltage clamp recordings were performed similar to neuronal recordings. Pyramidal neurons in layer 5 of the somatosensory cortex were selected based on their orientation in the slice and the visibility of processes especially the axon initial segment and the myelinated axon. During potassium application experiments, 50 ms long pulses with 140 mM K^+^ were pressure applied to the myelinated axon and the nodes of Ranvier. For these experiments, the slice was fixed with 4% PFA for subsequent immunohistochemistry labelling and confirmation of the morphology and the location of myelinated internodes.

### Pharmacology

Stock solutions of all pharmacological agents were prepared in distilled water or according to the datasheet provided by the manufacturer. For acute experiments drugs were then diluted in ACSF. Sodium channels were blocked with 0.5 µM tetrodotoxin (TTX, Alomone Labs, Israel). Kv1 channels were blocked with 100 nM Dendrotoxin-I (Alomone) and Kv7 channels were blocked with 20 µM XE991 (Alomone). Kir channels were blocked with 100 µM BaCl2 (Sigma-Aldrich). HCN channels were blocked with 30 µM ZD7288 (Sigma-Aldrich). TTX and BaCl2 were perfused for 5 minutes in the bath before recordings were started and all other blockers were perfused ≥14 minutes before recordings.

### Tissue fixation and immunohistochemistry

Half brains (**see Preparation of acute brain slices**) were dropped fixed in 4% PFA for 2 hours at room temperature and subsequently washed in PBS. Hemispheres were cryoprotected in 15% sucrose solution at 4°C until brains sank and then transferred and incubated at 4°C in 30% sucrose solution in PBS until the brains were saturated with the solution. Hemispheres were mounted and frozen onto the stage of a cryo-microtome (SM2010R, Leica) and coronal slices of 50 µm thickness were cut. Slices containing the ASAP3 injection site in the primary somatosensory cortex (limb region) were used for immunolabelling. Alternatively, 300 µm slices from acute experiments were fixed in 4% PFA and immunolabelling was performed after washing with PBS. To reduce unspecific immunolabeling, tissue was incubated in blocking buffer composed of 2.5% goat serum, 2.5% bovine serum albumin (BSA) and 0.3% triton X-100 in PBS at room temperature for 2 hours. Primary antibodies (Table 1) were diluted to their final concentration in blocking buffer and incubated at 4°C for 48 hours. For visualizing cell morphology with biocytin, we incubated tissue with streptavidin conjugated with Alexa-555 (1: 500, S21381, Invitrogen). After incubation with primary antibodies, slices were washed 3 times for 5 minutes in PBS and then incubated for 2 hours at RT with corresponding secondary antibodies diluted in PBS. After extensive washing, slices were mounted onto glass slides, air-dried for 10 minutes and covered by fluoroshield mounting medium (F6182, Sigma-Aldrich) and a coverslip.

**Table 1:** Antibodies used in this study.

| Antigen | Host | Dilution | Manufacturer | Identifiers (Order number, RRID) |
| --- | --- | --- | --- | --- |
| **Primary Antibodies** |  |  |  |  |
| Anti-GFP | Chicken polyclonal | 1:1000 and 1:200 for immuno EM | Aves Lab | Cat #: GFP-1020, RRID:AB_10000240 |
| Anti-CNPase, 11-5B, | Mouse monoclonal | 1:500 | Sigma-Aldrich | Cat #: C5922, RRID:AB_476854 |
| Anti-MBP | Mouse monoclonal | 1:1000 | Biolegend | Cat #: 808402, RRID:AB_2564742 |
| Anti-NeuN, A60 | Guinea pig polyclonal | 1:1000 | Sigma-Aldrich | Cat #: ABN90P, RRID:AB_2341095 |
| **Secondary Antibodies** |  |  |  |  |
| Goat polyclonal Alexa488 | anti-Chicken | 1:500 | Invitrogen | Cat# A-11039, RRID:AB_2534096 |
| Goat polyclonal Alexa568 | anti-Mouse IgG1 | 1:500 | Invitrogen | Cat#: A21124, RRID:AB_2535766 |
| Goat polyclonal Alexa647 | anti-Rabbit | 1:500 | Invitrogen | Cat#: A21245, RRID:AB_2535813 |
| Goat polyclonal Alexa647 | anti-Guinea pig | 1:500 | Invitrogen | Cat# A-21450, RRID:AB_141882 |
| Donkey anti IgY EM grade – 6 nm gold | Anti-chicken | 1:60 | Jackson Immuno | Cat# 703-195-155 RRID:AB_2340367 |
| **Streptavidins** |  |  |  |  |
| Streptavidin Alexa 555 |  | 1: 500 | Invitrogen | Cat# S21381 |

### Immunoelectron microscopy

Mice were perfused with room temperature PBS containing heparin (12.5 U/ml) followed by a freshly prepared fixation solution composed of 4% PFA and 0.25% glutaraldehyde (both Electron microscopy science) in PB. Brains were extracted and post-fixed for 2h in the same fixation solution. Vibratome sections of the brains were cut at 50 and 100 µm. Sections that contained the neocortical injection site were selected, photographed, and trimmed under an epifluorescence equipped stereomicroscope (blue excitation). Subsequently, trimmed tissue was submersed in 2.3 M sucrose solution in PB and stored at 4°C. The trimmed sections were then mounted and frozen on an aluminum pin before storage in liquid nitrogen. Further trimming under visual guidance was based on the photos and location of the expression location and was performed at −90°C with a cryotrimmer (trim20, Diatome Ltd, Nidau, Switzerland), mounted on an ultramicrotome (EM UC7 Leica) equipped with a cryochamber (FC7, Leica) Ultrathin 70 nm sections were cut at −120°C with a cryo-immuno 35° angled diamond knife (Diatome) and picked up with 2.3 M sucrose /1% methylcellulose (V:V) droplet, thawed and placed on nickel grids with 1% butvar film. For the immunohistological reaction, grids were washed 3 times with purified water for 2 minutes followed by a wash in PBS supplemented with 0.15% glycine for 15 minutes and then incubated with blocking solution for donkey gold conjugates (905.005, Aurion, Wageningen, The Netherlands) for 30 minutes. Subsequently, grids were incubated with primary antibody (anti-GFP, 1:200) diluted in PBS supplemented with 0.1% BSA-c (Aurion) for one hour at room temperature. We confirmed that the primary anti-GFP antibody is suitability to detect ASAP3 after fixation with PFA and glutaraldehyde in a separate set of experiments. After primary antibody incubation, grids were rinsed five times with PBS and 0.1% BSA-c, for five minutes. The secondary anti-chicken antibody was coupled with a 6 nm gold particle (1:60 dilution, Jackson Immuno) and was diluted in PBS supplemented with 0.1% BSA-c and incubated for one hour. Afterwards, samples were washed five times with PBS supplemented with 0.1% BSA-c for five minutes. A second wash was performed two times with PBS for two minutes before the antibody labelling was stabilized with 1% glutaraldehyde in PBS for five minutes. Afterwards, the grids were rinsed three times with PBS for two minutes followed by five rinses with double-distilled water for one minute. As negative control we used the same tissue sections, but omitted the primary antibody. Control sections were otherwise treated the same. As a final step, grids were floated in drops of ice-cold 0.4% uranyl acetate, 1.8% methylcellulose, pH 4, in distilled water for 10 minutes, in the dark and collected with a wire loop. Images of section were taken on a Hitachi H7650 TEM equipped with a SC1000 ORIUS CCD camera (Gatan, Pleasanton, CA, USA) at varying magnifications.

### Confocal microscopy

Confocal images were acquired with Leica TCS SP5 confocal microscopes (Leica Microsystems, Germany). Overview images were acquired with a 10x 0.3NA objective (HCX PL Fluotar, Leica). Tiled z-stack were acquired with a 40× 1.30NA objective (HC PL APO CS2 OIL UV, Leica) or a 63× 1.40NA objective (HCX PL APO CS OIL UV, Leica). Individual cells or myelin sheaths were acquired using a 63× 1.40 NA objective and for some images we applied a digital zoom (from 2 to 4.5).

### Quantification and analysis

#### Image analysis of optical voltage changes

Image stacks of optically recorded voltage changes in oligodendrocytes were analyzed in FIJI [(Schindelin et al., 2012), RRID:SCR_002285], Excel (RRID:SCR_016137) and AxoGraph (RRID: SCR_014284, AxoGraph Scientific, Sydney, Australia). After selecting ROIs for the target (cell body or myelin sheath) and the background that contained no visible process, the raw fluorescence signals were extracted from FIJI. Data from each acquisition/stack was then pre-processed using an Excel template. First, an average background was determined during baseline recordings that was then subtracted from all other ROIs in the recording. From the background subtracted signal ΔF/F0 was calculated by dividing the entire signal with an average baseline fluorescence (F0) that was calculated as average between –60 to –10 ms before the onset of stimulation. For 2-photon data, pre-processing of images including the extraction of ROI, background and calculation of ΔF/F0 was performed with Femtonics software (mES). Resulting ΔF/F0 traces were plotted in Axograph to determine peak amplitude, the area under the curve and decay times of the signal (Exponential fit function). Voltage traces were inverted for display to reflect the depolarization of the oligodendrocyte during the experiment and we report the optical voltage change as –ΔF/F0.

#### Immunogold EM analysis

Immunogold labelling was quantified from images we obtained from 3 mice and a minimum of 17 images per mouse with an average of 315 ± 30 gold particles per mouse across all analyzed images. On average the analyzed images covered 61 ± 29 µm^2^. Gold particles were assigned to different cellular structures that were identified and included myelin (compact, inner, outer), axon, mitochondria, other and no identifiable structure. No gold particles were detected in the negative controls (n = 3 mice).

#### Statistical analysis

Statistical analysis was done with Prism 10 (GraphPad Software, Boston, MA, USA, RRID:SCR_002798). Normal distribution of the data was tested with a Shapiro-Wilk test. For the comparison between two paired groups without normally distributed data or with n ≤ 6, a Wilcoxon test was used. Two paired groups with n > 6 and normally distributed data were tested with a paired t-test.

In the case of two non-paired groups without normal distribution or n ≤ 6, a Mann-Whitney test was applied. For normally distributed data of two non-paired groups and n > 6 a student t-test was used. In the case of more than two groups, paired, without normal distribution, the Friedman test was applied. For more than two groups non-paired and non-normally distributed, the Kruskal-Wallis test was performed.

The data were considered to be statistically significant when the p-value < 0.05. The data shown in the figures or presented in the manuscript are given as mean ± standard error of the mean (S.E.M.).

## Acknowledgments

We thank Saroop Sanghera for experimental support, Nicolas Snaidero for helpful discussion and Sebastian Marais for technical support for the 2P microscope. We thank the PIV-EXPE of the University of Bordeaux for animal care and the genotyping facility of Neurocentre Magendie for genotyping of experimental animals.

## Competing interests

The authors declare no conflict of interest.

### Author contributions

Conceptualization: AB; Investigation: ML, MP, AB; Formal analysis: ML, MP, AB. Funding acquisition: AB; Writing – original draft: AB, Writing – review & editing: ML, MP, AB.

---

**Supplementary Figure 1:**
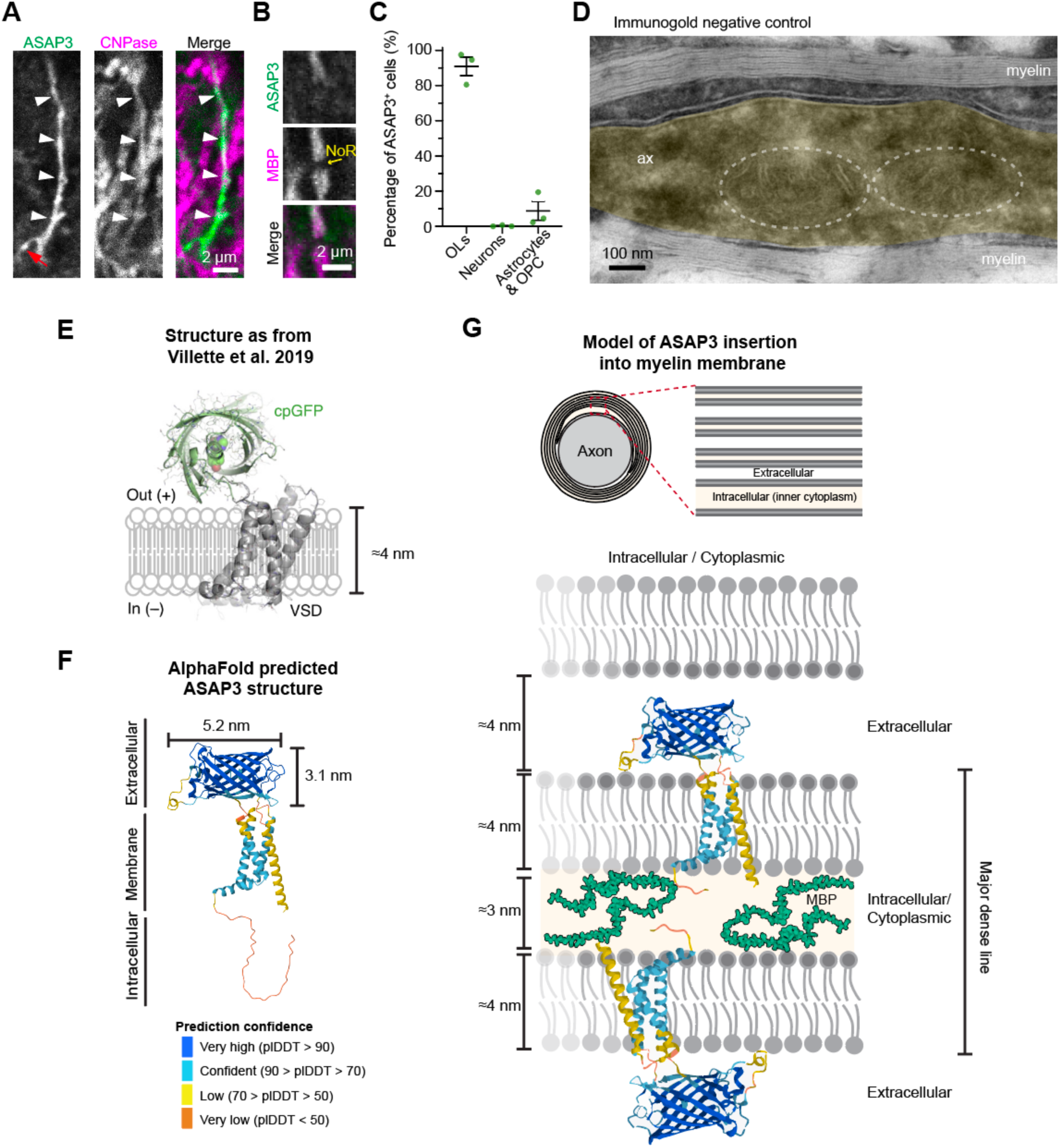
Examples and quantification of ASAP3 expression in oligodendrocytes. (A) Confocal images of ASAP3 expressing oligodendrocytes co-labelled with CNPase. White arrowheads point to an ASAP3 positive myelin sheath and the red arrow to a connecting process. (B) Higher magnification images of an ASAP and MBP positive myelin sheath that ends at a Node or Ranvier (NoR) and is flanked by a MBP positive ASAP3^–^ myelin sheath. (C) Quantification of ASAP3 cell specificity shows that the majority of ASAP3^+^ cells are oligodendrocytes (CNPase or Olig2 staining and morphology) and about 10% of cells were identified as astrocytes or OPCs based on their morphology and absence of CNPase labelling. (D) Transmission electron microscopy image after omission of the primary antibody shows the absence of immunogold particles in an axon covered by myelin. The axon is pseudo colored in yellow and mitochondria in the axon are encircled with dashed lines. (E) 3D model of ASAP3 as previously published (Vilette et al, 2019). (F) Alpha-fold model prediction of ASAP3 structure is comparable to the previously published model. The structure of the N-terminus could not be predicted with high accuracy. The first 41 amino acids were predicted to be located in the cytoplasm but the prediction differs from the arrangement in A. (G) Schematic model of compact myelin and the insertion of ASAP3 into the myelin membrane. Based on the ASAP3 design, the cpGFP domain locates into the extracellular space, whereas the voltage sensing domain is inserted within the membrane and only a small part is within the cytoplasm. While immuno EM demonstrated the location within compact myelin, it remains unresolved whether ASAP3 is able to diffuse laterally within compact myelin after integration.

**Supplementary Figure 2:**
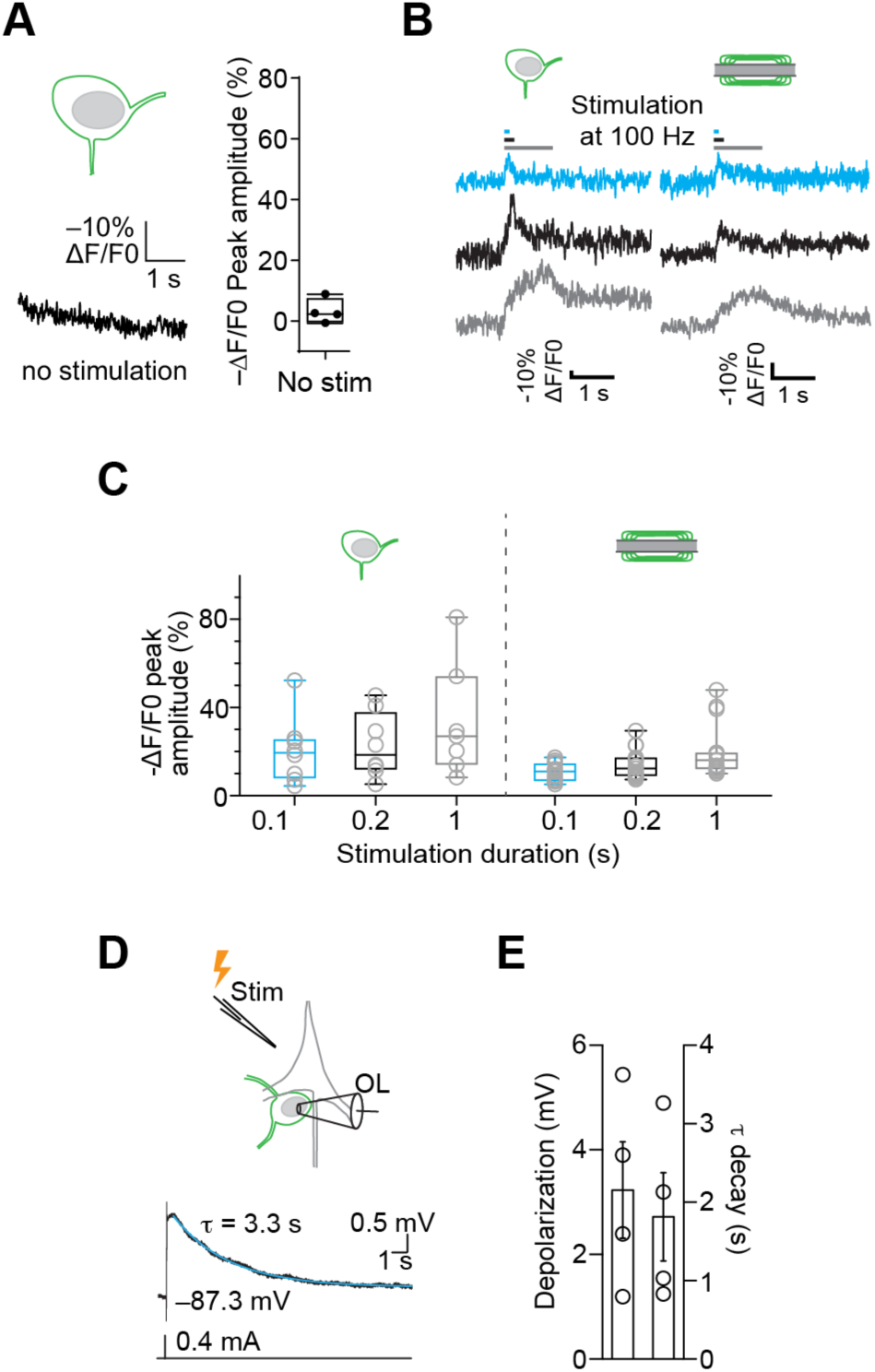
Optical membrane voltage measurements in oligodendrocytes. (A) Voltage imaging trace of a cell body without stimulation shows the absence of optically recorded voltage changes. Summary plot of several experiments in the absence of stimulation (n = 4 cells). (B) Voltage changes vary with stimulation lengths from 0.1 to 1 second. One second stimulation shows a non-linearity stemming from reduction of neuronal firing as a consequence of a depolarization block towards the end of the stimulus. (C) Quantification and box plots of the peak optically measured voltage change (ΔF/F0) for the different stimulation lengths for oligodendrocyte cell bodies and myelin sheaths. Cell body: n = 8 cells from 4 animals; myelin n = 14 myelin sheaths from 3 animals. (D) *Top:* Schematic of whole-cell recording and simultaneous extracellular stimulation of a single oligodendrocyte. *Bottom:* A single extracellular stimulation elicited a somatic depolarization that decayed slowly in a current-clamped oligodendrocyte located in satellite position (sOL) to a neuron. (E) Summary plot displaying peak depolarization and the decay time constant of a single extracellular stimulation (n = 4 oligodendrocytes from 2 mice).

**Supplementary Figure 3:**
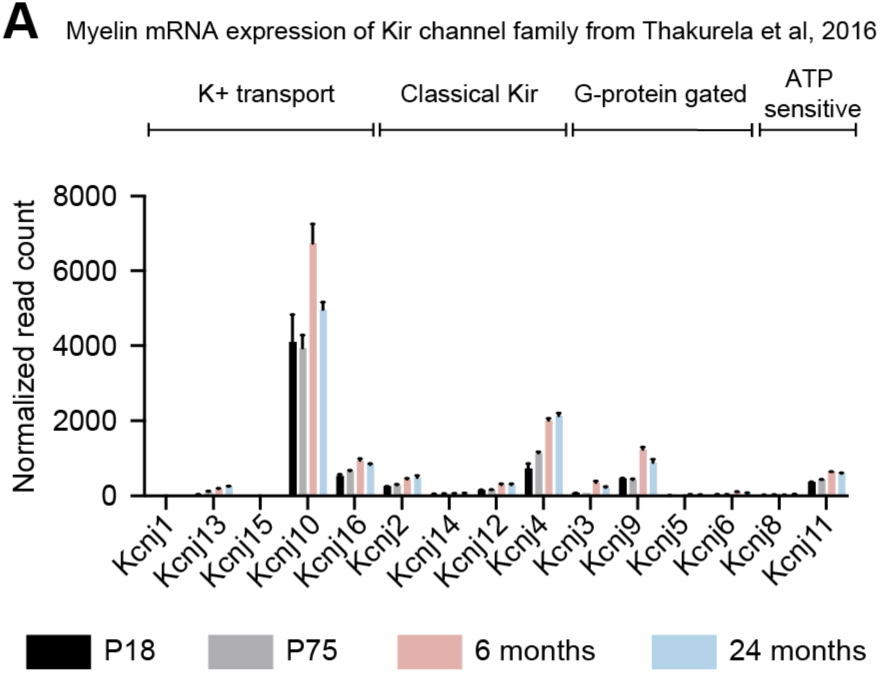
mRNA expression of Kir channels in myelin. (A) mRNA expression data showing the normalized read count for the KCNJ (Kir channel) family. Genes were grouped by function (top) and for four different ages. Only Kir4.1 corresponding to KCNJ10 was also detected on the protein level by mass spectrometry. Data adapted from Thakurela et al (2016).

