## Supplementary Figures for "Neuronal activity induces myelin voltage changes that reflect action potential dependent myelin potassium buffering"

This file contains 3 supplementary Figures

### Supplementary Figure 1

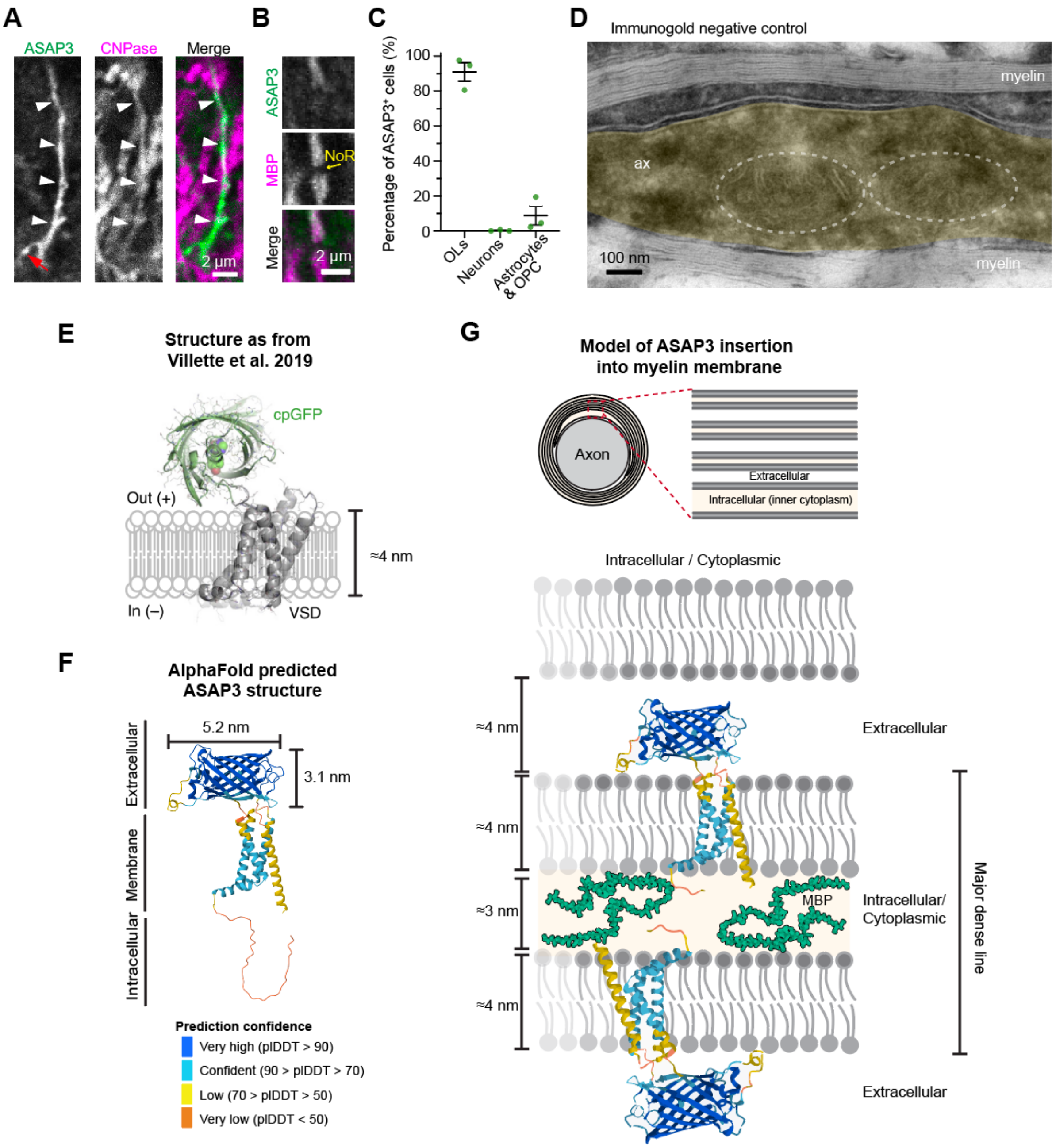


**Supplementary Figure 1: Examples and quantification of ASAP3 expression in oligodendrocytes**

(A) Confocal images of ASAP3 expressing oligodendrocytes co-labelled with CNPase. White arrowheads point to an ASAP3 positive myelin sheath and the red arrow to a connecting process.

(B) Higher magnification images of an ASAP and MBP positive myelin sheath that ends at a Node or Ranvier (NoR) and is flanked by a MBP positive ASAP3$^{-}$ myelin sheath.

(C) Quantification of ASAP3 cell specificity shows that the majority of ASAP3$^+$ cells are oligodendrocytes (CNPase or Olig2 staining and morphology) and about 10% of cells were identified as astrocytes or OPCs based on their morphology and absence of CNPase labelling.

(D) Transmission electron microscopy image after omission of the primary antibody shows the absence of immunogold particles in an axon covered by myelin. The axon is pseudo colored in yellow and mitochondria in the axon are encircled with dashed lines.

(E) 3D model of ASAP3 as previously published (Vilette et al, 2019).

(F) Alpha-fold model prediction of ASAP3 structure is comparable to the previously published model. The structure of the N-terminus could not be predicted with high accuracy. The first 41 amino acids were predicted to be located in the cytoplasm but the prediction differs from the arrangement in A.

(G) Schematic model of compact myelin and the insertion of ASAP3 into the myelin membrane. Based on the ASAP3 design, the cpGFP domain locates into the extracellular space, whereas the voltage sensing domain is inserted within the membrane and only a small part is within the cytoplasm. While immuno EM demonstrated the location within compact myelin, it remains unresolved whether ASAP3 is able to diffuse laterally within compact myelin after integration.

### Supplementary Figure 2

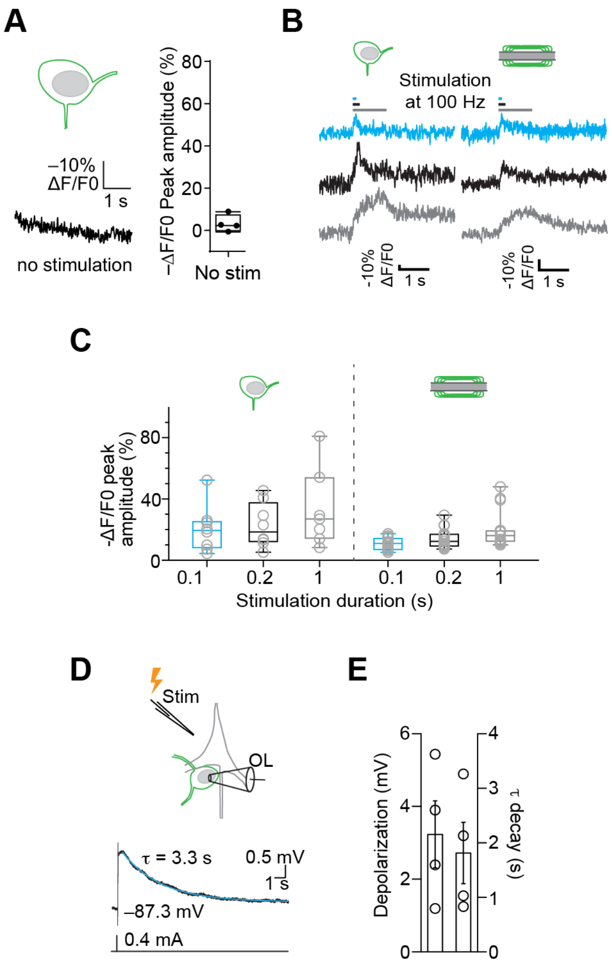


**Supplementary Figure 2: Optical membrane voltage measurements in oligodendrocytes**

(A) Voltage imaging trace of a cell body without stimulation shows the absence of optically recorded voltage changes. Summary plot of several experiments in the absence of stimulation (n = 4 cells).

(B) Voltage changes vary with stimulation lengths from 0.1 to 1 second. One second stimulation shows a non-linearity stemming from reduction of neuronal firing as a consequence of a depolarization block towards the end of the stimulus.

(C) Quantification and box plots of the peak optically measured voltage change (ΔF/F0) for the different stimulation lengths for oligodendrocyte cell bodies and myelin sheaths. Cell body: n = 8 cells from 4 animals; myelin n = 14 myelin sheaths from 3 animals.

(D) *Top:* Schematic of whole-cell recording and simultaneous extracellular stimulation of a single oligodendrocyte. *Bottom:* A single extracellular stimulation elicited a somatic depolarization that decayed slowly in a current-clamped oligodendrocyte located in satellite position (sOL) to a neuron.

(E) Summary plot displaying peak depolarization and the decay time constant of a single extracellular stimulation (n = 4 oligodendrocytes from 2 mice).

### Supplementary Figure 3

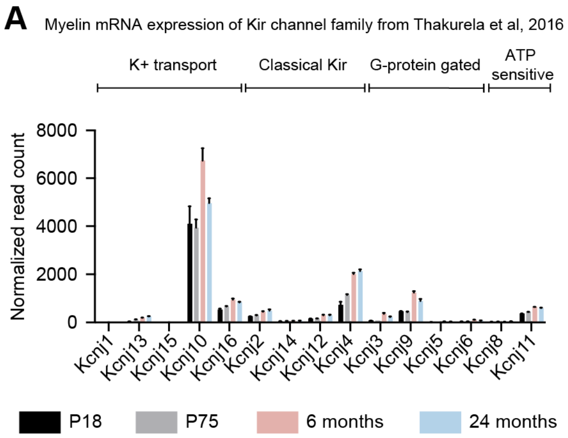


**Supplementary Figure 3: mRNA expression of Kir channels in myelin**

(A) mRNA expression data showing the normalized read count for the KCNJ (Kir channel) family. Genes were grouped by function (top) and for four different ages. Only Kir4.1 corresponding to KCNJ10 was also detected on the protein level by mass spectrometry. Data adapted from Thakurela et al (2016).
